## Supplementary material for "Mapping quantitative trait loci and predicting candidate genes for leaf angle in maize": S1 Table RT-qPCR primer

| Primer name | Primer sequence(5' to 3') |
| --- | --- |
| qRTzm5682-F | GGTCACTCGACCAGCTAAAT |
| qRTzm5682-R | CCTCCCTTGCCGATGAGGTT |
| qRTzm5808-F | GCAAGGAATTGGTCGAAGGG |
| qRTzm5808-R | TCGCCTGCAATTCCCTTTTC |
| qRTzm5818-F | AATCAGGTCATCGCAGAGGT |
| qRTzm5818-R | TTGCCTAACGATCTCTCCTC |
| qRTzm5823-F | AGCATGGAGGAGATCTTCGG |
| qRTzm5823-R | ACGCACTCTGGACAAACAAC |
| qRTzm6036-F | TTGCTGGATGTCACACCTCT |
| qRTzm6036-R | GTAGCCATCTCCCTCTCACC |
| qRTzm6153-F | GTCAACATTGGCTCTGCTCC |
| qRTzm6153-R | ATCACCCTAAAACGGCTTGC |
| qRTzm6258-F | GCAGATGTCACAGGAGGAGT |
| qRTzm6258-R | AACAACGATCTGATGCTGCC |
| qRTzm6296-F | GCTGTTATTGAGGCGCTGAA |
| qRTzm6296-R | TTGCTCATACTGCTGGGGAA |
| qRTzm6443-F | ACACAGGCAAAAGCTTCAGG |
| qRTzm6443-R | ACAATTGCCCGCCACATATC |
| qRTzm6467-F | CATCAAGTACCCGTCGGAGA |
| qRTzm6467-R | CTAGCATCCCAGTTCCCTCC |
| qRTzm6629-F | AGTACATTGCTCACAGGCCT |
| qRTzm6629-R | TAATCTAGGGCGAGGCTTGG |
| qRTzm6631-F | TCGCTTTGTTGTTGGTCTGG |
| qRTzm6631-R | GAGGACAATGCAAGCAACCA |
| qRTzm6646-F | CGACTTCCTCTTCGCCTTCT |
| qRTzm6646-R | ATTCCTCGTCGGCAATCAGT |
| qRTzm6688-F | GGGTGGAATGATTGTCTCGC |
| qRTzm6688-R | TCTCCGAGTTGTACACACCC |
| qRTzm6700-F | GACACATGGCCTGCTTTCAA |
| qRTzm6700-R | GAACATCGAAGGAGCAGCAG |
| qRTzm9580-F | CACCCGAAATCAACGAGCAT |
| qRTzm9580-R | GGAGTGGGAGTGGTGATGAT |
| qRTzm9593-F | AGGTATGGGGCTAGGATTGC |
| qRTzm9593-R | AAGTCCAGAGCCAGTACCAC |
| qRTzm9594-F | GGATCCTGGGCGATGTTTTC |
| qRTzm9594-R | CCGGCCGCATAAATACACAA |
| qRTzm9610-F | GCCTACTACCTACTCAAAGCCT |
| qRTzm9610-R | GGAAACGAACACCTGGTATCTG |
| qRTzm9611-F | CAACTGCTTGACCTGGACG |
| qRTzm9611-R | CTAGTTGGGCCTCAGGTTGT |
| qRTzm9679-F | CTTGGCAGAGTGGTAAGGGA |
| qRTzm9679-R | ACTCAAGCAGTCCCCTTCTC |
| qRTzm9813-F | GATCTTTCGGGGCATTCGTC |
| qRTzm9813-R | TGTCAAGGTAGTGGCGAAGT |
| qRTzm9835-F | GACCCTTGAGCAGAGAGAGG |
| qRTzm9835-R | CATGAACCTCCTGAGCCTGA |
| qRTzm10000-F | TCATCAGGGCTTTCTCGTGT |
| qRTzm10000-R | CGGAGAAGCTACTGGTGACT |
| qRTzm10009-F | GTGTGGGAAGGAGACGAGAA |
| qRTzm10009-R | TGTCTCACGGGTGTTCTTGA |
| qRTzm10152-F | AGTGGAGATGCTGGGTATGAAT |
| qRTzm10152-R | TCACCTTATCATCGCCCACATA |
